# Acetylsalicylic acid and Salicylic acid inhibit SARS-CoV-2 replication in precision-cut lung slices

**DOI:** 10.1101/2022.08.09.503270

**Authors:** Nina Geiger, Eva-Maria König, Heike Oberwinkler, Valeria Roll, Viktoria Diesendorf, Sofie Fähr, Helena Obernolte, Katherina Seewald, Sabine Wronski, Maria Steinke, Jochen Bodem

## Abstract

Aspirin, with its active compound acetylsalicylic acid (ASA), shows antiviral activity against rhino- and influenza viruses and at high concentrations. We sought to investigate whether ASA and its metabolite salicylic acid (SA) inhibit SARS-CoV-2 since it might use similar pathways to influenza viruses. The compound-treated cells were infected with SARS-CoV-2. Viral replication was analyzed by RTqPCR. The compounds suppressed SARS-CoV-2 replication in cell culture cells and in a patient-near replication system using human precision-cut lung slices by two orders of magnitude. The compounds did not interfere with viral entry but led to lower viral RNA expression after 24 h.

---

Starting in December 2019, SARS-CoV-2 became a pandemic thread with more than 580 million cases worldwide and more than 6.35 million deaths until the summer of 2022, with still rising numbers due to new, more efficiently transmitted variants of SARS-CoV-2 and lower prevention measures^1^. In contrast to previous years, this combination further increased infection rates in Western countries even during the summer months^1^. The high infection rates compensated in Germany in parts the effects of the introduction of efficient vaccines leading to death rates comparable to April 2020 at the beginning of the pandemic, when no vaccine had been available. Unfortunately, efficient direct-acting therapies are still widely unavailable and have severe adverse side effects, such as the potential teratogenicity of Molnupiravir^2^.

Several approaches for off-label use of approved drugs against SARS-CoV-2, such as Remdesivir^3,4^, Fluoxetine^5^ and Fluvoxamine^6^, were published, and some were incorporated into the national treatment guidelines, but others, for instance, Chloroquine, failed in patients due to inadequate preclinical test systems^7^.

SARS-CoV-2 is a plus-stranded, enveloped RNA virus. It uses the spike protein to enter the cells via angiotensin-converting enzyme 2 (ACE2), which is abundantly expressed in alveolar and heart tissue. The virus enters the cell via cell type-specific pH-dependent endocytosis or by direct fusion with the cytoplasmic membrane depending on the concentration of the cellular TMPRSS2 proteases^8^. Inhibition of viral entry will result in lower RNA amounts early after infection. The incoming RNA is directly translated into viral proteins and expressed by the viral RNA polymerase. Thus, inhibition of viral replication steps after entry until RNA expression will result in a lower increase of RNA amounts in the cells.

Cell culture-based infection experiments were conducted to determine the potential effects of ASA and SA on the SARS-CoV-2 replication. First, the cytotoxicity of the compounds was analyzed, and secondly, the impact on viral replication rates was measured. A549-ACE2, Huh-7 and Vero cells were incubated with ASA or SA for three days to investigate cytotoxicity, and the relative cell growth was determined by automatic cell counting as described before^5^. Concentrations of 3 mM SA or ASA did not influence cell growth.

Since it has been shown that the antiviral effect of a defined compound on SARS-CoV-2 replication depends on the applied test system^9^ the influence of ASA and SA on viral replication in Vero, A549-ACE2 and Huh-7 cells was compared. The cells were pre-incubated with the compounds at 1.5 and 3 mM and infected with SARS-CoV-2 in triplicates (Figure 1). All infections were repeated twice.

**Figure 1:**
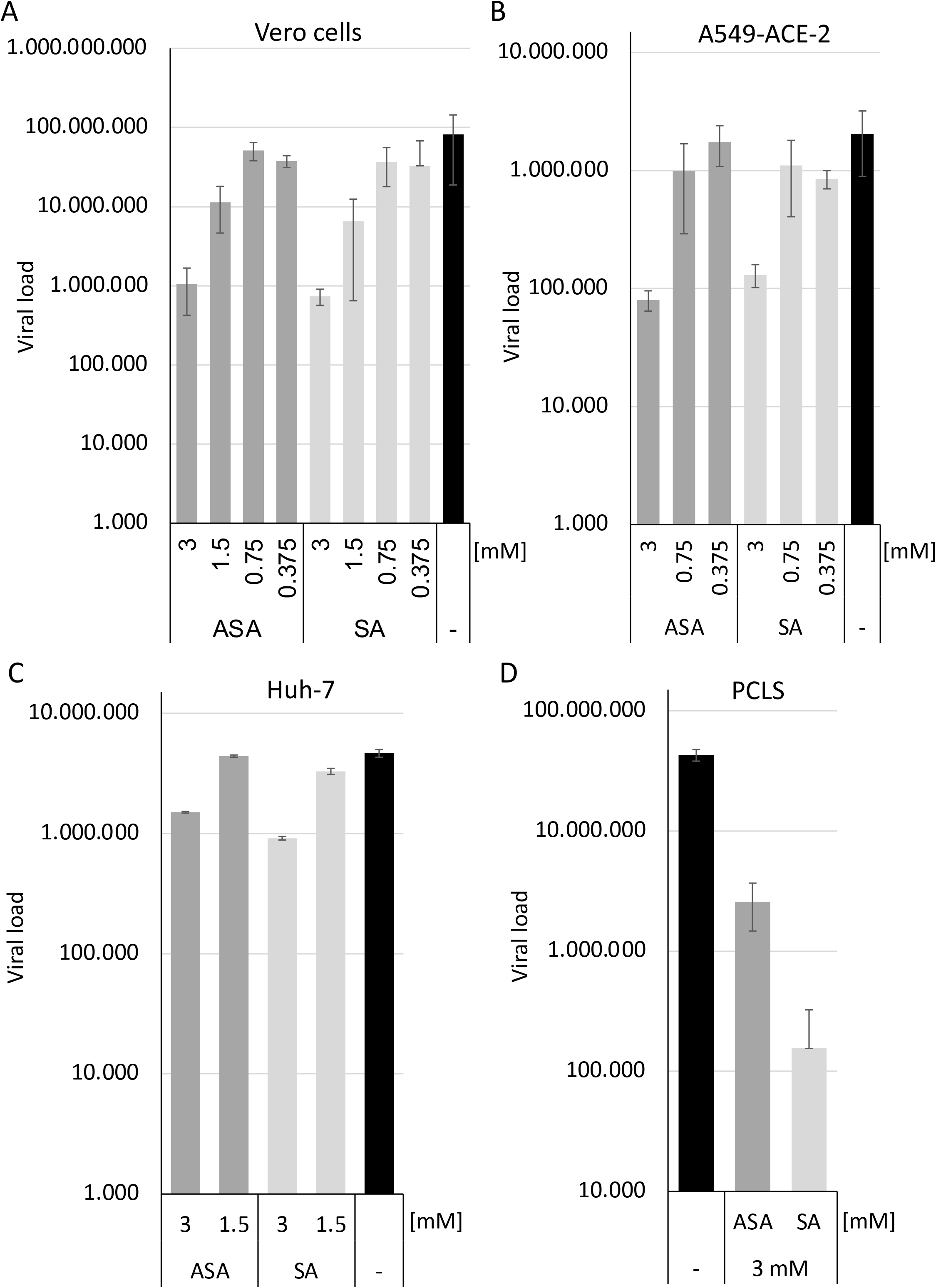
SA and ASA inhibit viral replication in Vero, A549-ACE2 and Huh-7 cells and human precision-cut lung slices. Vero (A), A549-ACE2 (B) and Huh-7 (C) cells, as well as PCLS, were treated with the compounds and infected with SARS-CoV-2. Viral loads were determined 3 days after infection. Bars represent the mean, and error bars the standard deviation.

Cellular supernatants were collected two days after infection. Viral RNAs were extracted using a MagNA Pure 24 system (Roche, Mannheim). SARS-CoV-2 RNA genomes were quantified with the dual-target SARS-CoV-2 RdRP RTqPCR assay kit with RNA Process Control PCR Kit (Roche) with a LightCycler 480 II (Roche) (Figure 1). ASA and SA significantly inhibited viral replication in a cell type-specific way by approximately half (Huh-7) or two (Vero cells and A459-ACE2) orders of magnitude, respectively (Figure 1A/B/C). This reduction is above the previously reported suppression of the respiratory syncytial, Influenza, and Rhinoviruses^10^. It indicates that both ASA and SA interfere with viral replication, indicating that the compounds might inhibit an essential SARS-CoV-2 pathway.

Recent studies with antiviral drugs have revealed that the translation from a cell culture-based system, such as VeroE6 cells might result in inactive patient therapies. Chloroquine, for example, shows antiviral activity in these cells but neither in Calu3 cells, human epithelial tissue models, macaques or in patients^7,9^. Therefore, we decided to determine the effects of ASA and SA in human precision-cut lung slices (PCLS) as described before^5^. The PCLS were incubated for 1 h at 37°C, incubated with the compounds and infected with SARS-CoV-2 at a high MOI of 10. After 3 days, 10 μl of the cell culture supernatants were used to infect Vero cells to determine the infectious viruses. Cellular supernatants were collected, RNAs isolated and viral genomes quantified after 3 days by RTqPCR (Figure 1D). ASA inhibited viral replication by approximately one order of magnitude, while with SA, a reduction of more than two orders of magnitudes was achieved. The treatment with the compounds reduces viral replication in all cell lines analysed and in PCLS, indicating that they are promising candidates to be evaluated in antiviral therapy.

To determine whether ASA and SA inhibit the entry of SARS-CoV-2, A549-ACE2 cells were incubated with the substances, and viral RNA expression was determined 6 h and 24 h after infection. All infections were performed in independent triplicates. After one hour, the medium exchanged to remove the virus from the supernatant. After 6 h, the cells were washed with ice-cold PBS and the RNAs were prepared according to the manufacturer’s instructions (Omega, Germany). Similar RNA amounts of (150 ng) were used in RTqPCR reactions to quantify SARS-CoV-2 RNA as described above. Results were normalized on GAPDH by RTqPCR. After 6 h of infection, we observed similar SARS-CoV-2 RNA amounts in all lysates (Figure 2A) providing evidence that equal amounts of the virus could enter the cells despite the treatment. Thus, we conclude that the viral entry process is not targeted in SARS-CoV-2 infections. The low amount of viral RNAs at 6 h indicates that this time-point is before substantial viral gene expression.

**Figure 2:**
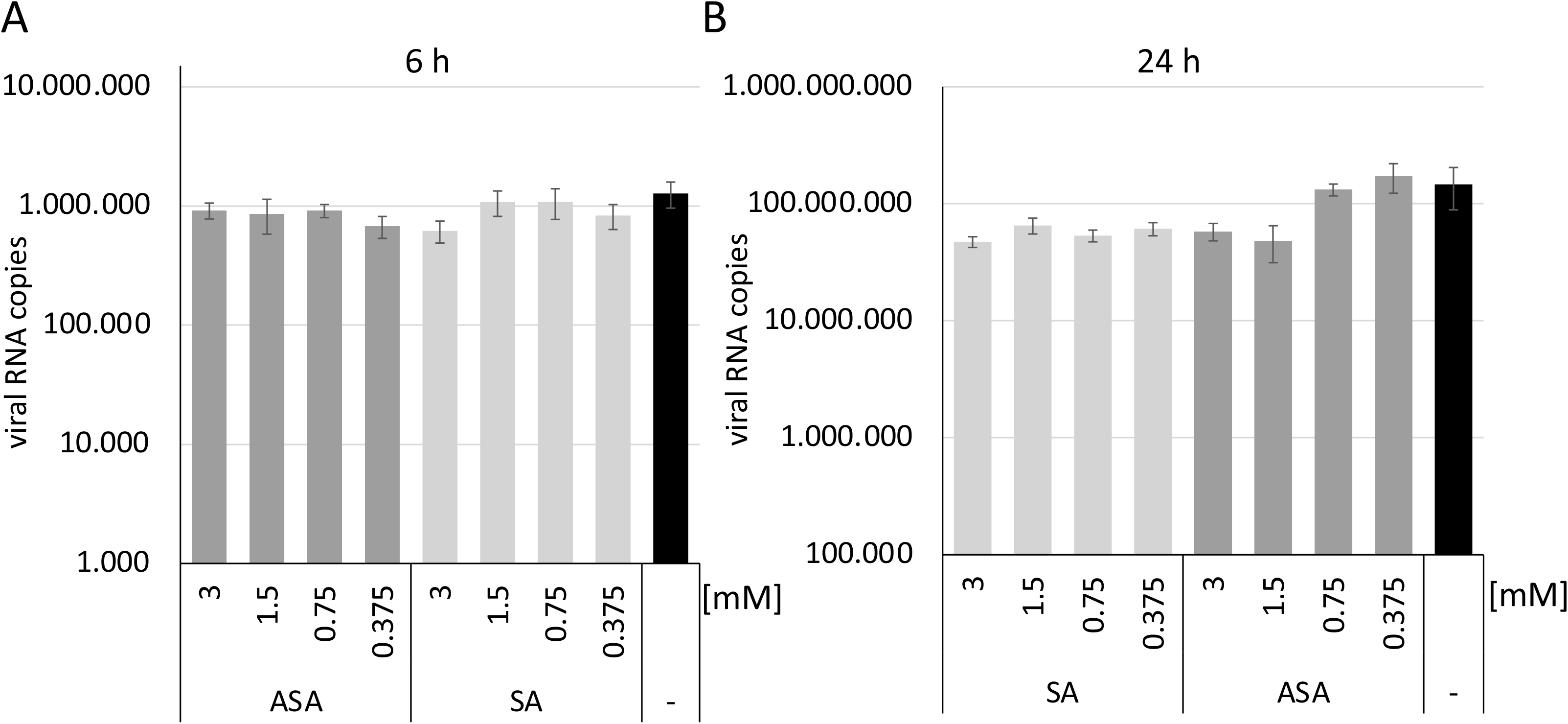
SA and ASA do not interfere with viral entry but with replication steps before or during gene expression. Total RNAs of SARS-CoV-2 infected cells were isolated after 6 and 24 h, and SARS-CoV-2 RNA was quantified by RTqPCR. Bars represent the mean, and error bars the standard deviation.

Similar experiments were performed to analyze the infection after 24 h in the presence of both compounds (Figure 2B). We observed a reduction of viral RNA amounts after 24 h with ASA at concentrations higher than 1.5 mM. In comparison, SA reduced viral RNAs at lower concentrations, similar to the results shown above. This indicates that the compound inhibits replication steps before or during gene expression.

In summary, ASA and SA are effective against SARS-CoV-2 in different animal- and human-derived cell lines and in the precision-cut, patient-derived lung slices.

## Acknowledgements

Bayer Vital GmbH funded a part of this study.

## Ethics statement

Human lung lobes were acquired from patients undergoing lobe resection for cancer at Hannover Medical School. The use of the tissue for research was approved by the ethics committee of the Hannover Medical School and complies with the Code of Ethics of the World Medical Association (number 2701–2015). All experiments were performed by relevant guidelines and regulations. All patients gave written informed consent to use explanted lung tissue for research and publish the results. No sample tissues were procured from prisoners.

